# Chloride intracellular channel 4 (CLIC4) is a global regulator of type 1 interferon signaling in Systemic Sclerosis (SSc) epithelial cells

**DOI:** 10.1101/2024.03.08.583925

**Authors:** Christopher W Wasson, Sophie L Dibb, Begoña Caballero-Ruiz, Eva M Clavane, Rebecca Wells, Vishal Kakkar, Enrico De Lorenzis, Rebecca L Ross, Jessica Bryon, Emma Derrett-Smith, Christopher P Denton, Paul J Meakin, Francesco Del Galdo

## Abstract

**Objectives:** Systemic sclerosis (SSc) is an autoimmune disease in which an immune-related injury induces fibrosis of the skin, progressing to affect the internal organs in the most serve cases. Type 1 interferon (IFN) signaling plays a major role in SSc disease progression. We have previously shown the chloride intracellular channel 4 (CLIC4) is upregulated in SSc skin fibroblasts and plays an important role in SSc fibrosis. In this study we investigated the role of CLIC4 in SSc keratinocyte biology.

**Methods:** healthy (HC) and SSc skin biopsies were analysed by immunohistochemistry for the expression of CLIC4. The skin keratinocyte cell line Hacats was stimulated with a range of type 1 IFN signaling agonists (POLY I:C, POLY dA:Dt, ODN 2216 and IFN-α). CLIC4 was inhibited with the chloride channel inhibitors NPPB and IAA-94 or siRNA. Conditioned media from HC or SSc fibroblasts was employed for indirect co-culture of Hacats and HUVECs.

**Results:** SSc skin biopsies showed high levels of CLIC4 in SSc skin fibroblasts, keratinocytes and endothelial cells compared to HC. Co-culture of Hacats and Huvecs with SSc fibroblast media induced CLIC4 expression. CLIC4 played an important role in type 1 IFN signalling in keratinocytes. Inhibition of CLIC4 blocked TLR3, TLR9 and cGAS mediated activation of the type 1 IFN signaling pathway. Additionally, inhibition of CLIC4 prevented SSc fibroblast media from inducing a type 1 IFN response in keratinocytes.

**Conclusions:** The data presented in this study suggests CLIC4 is a global regulator of type 1 IFN signalling in SSc epithelial cells.

**Key Messages:** *What is already known:* SSc disease progression is driven in part by a Type 1 IFN signature and CLIC4 has previously been implicated in SSc fibroblast activation.

*What this study adds:* We show for the first time CLIC4 is a regulator of type 1 interferon signalling in epithelial cells and plays an important role in the signalling found in SSc skin.

*How this study might affect research:* Targeting CLIC4 in the context of SSc may disrupt the fibrosis and inflammation associated with SSc.

## Introduction

Systemic Sclerosis (SSc) is an auto-immune disease which initially presents in the hands and feet of patients and can variably progress to affect most internal organs including the gastrointestinal tract, heart and lungs. The tissue fibrosis is associated with a deregulated activation of type-1 interferon (IFN) signalling (1,2) which leads (through activation of cytokines such IL-6 and TGF-β) to activation of myofibroblasts. Recent work from our group has shown the chloride intracellular channel 4 (CLIC4) is a mediator of SSc myofibroblast activation (3). Channel expression is increased in SSc dermal fibroblasts and inhibition of the channel with small molecule inhibitors blocks SSc myofibroblast activation (3). CLIC4 expression is regulated in dermal fibroblasts by the TGF-β, Wnt-3a and hedgehog signalling pathways (4). In addition, there is evidence that CLIC4 can activate the TGF-β signalling pathway by maintaining SMAD3 phosphorylation (5).

Keratinocytes and endothelial cells have been shown to contribute to SSc disease progression in the skin. The keratinocytes are a major source of type-1 IFN signalling in SSc skin (6) and angiogenesis is disrupted in SSc endothelial cells which contributes to the vascular involvement in SSc (7). Interestingly there is evidence that CLIC4 can modulate keratinocyte and endothelial cell biology. Overexpression of CLIC4 in keratinocytes leads to reduced mitochondrial membrane potential and the release of cytochrome C resulting in apoptosis (8). Furthermore stress inducers such as actinomycin D and etoposide can drive CLIC4 nuclear localization in keratinocytes (9). CLIC4 knockdown can prevent Ca(2+)-induced keratinocyte differentiation (10). In the context of endothelial cells, CLIC4 is important for endothelial cell proliferation (11), can mediate G protein coupled receptor signaling associated with vascular development (12) and has important roles in regulating endothelial cell apoptosis (13). Finally aberrant expression of CLIC4 is associated with idiopathic pulmonary hypertension (IPH) as inhibition of CLIC4 in IPH patient derived endothelial cells restores angiogenesis (14).

In this study, we aim to investigate the role of CLIC4 in SSc keratinocyte and endothelial cell biology. We show for the first time that CLIC4 is highly expressed in keratinocytes and endothelial cells of SSc patient skin and this is driven in part by TGF-β. CLIC4 is shown to be important for regulating STAT1 activation in the context of a type-1 IFN response in SSc keratinocytes.

## Materials and Methods

Materials and Methods can be found in supplementary materials

## Results

### CLIC4 levels are elevated in SSc skin keratinocytes, fibroblasts and endothelial cells

Previously we have shown CLIC4 expression levels were increased in dermal fibroblasts isolated from SSc patient skin (3,4) but the expression profile in SSc skin remained unknown. We analysed CLIC4 levels in SSc skin biopsies by immunohistochemistry (Figure1A, Supplementary Figure1A). We found CLIC4 levels were increased in dermal fibroblasts of the SSc skin compared to healthy control validating our previous findings. Interestingly we found high levels of CLIC4 in the keratinocytes and endothelial cells in the SSc skin biopsies (Figure1A). CLIC4 levels were high in the lower layers of the epidermis in SSc skin (Figure1A). Further analysis of isolated SSc keratinocytes found CLIC4 transcript levels were increased (1.5-fold) compared to healthy control (Figure1B).

**Figure 1:**
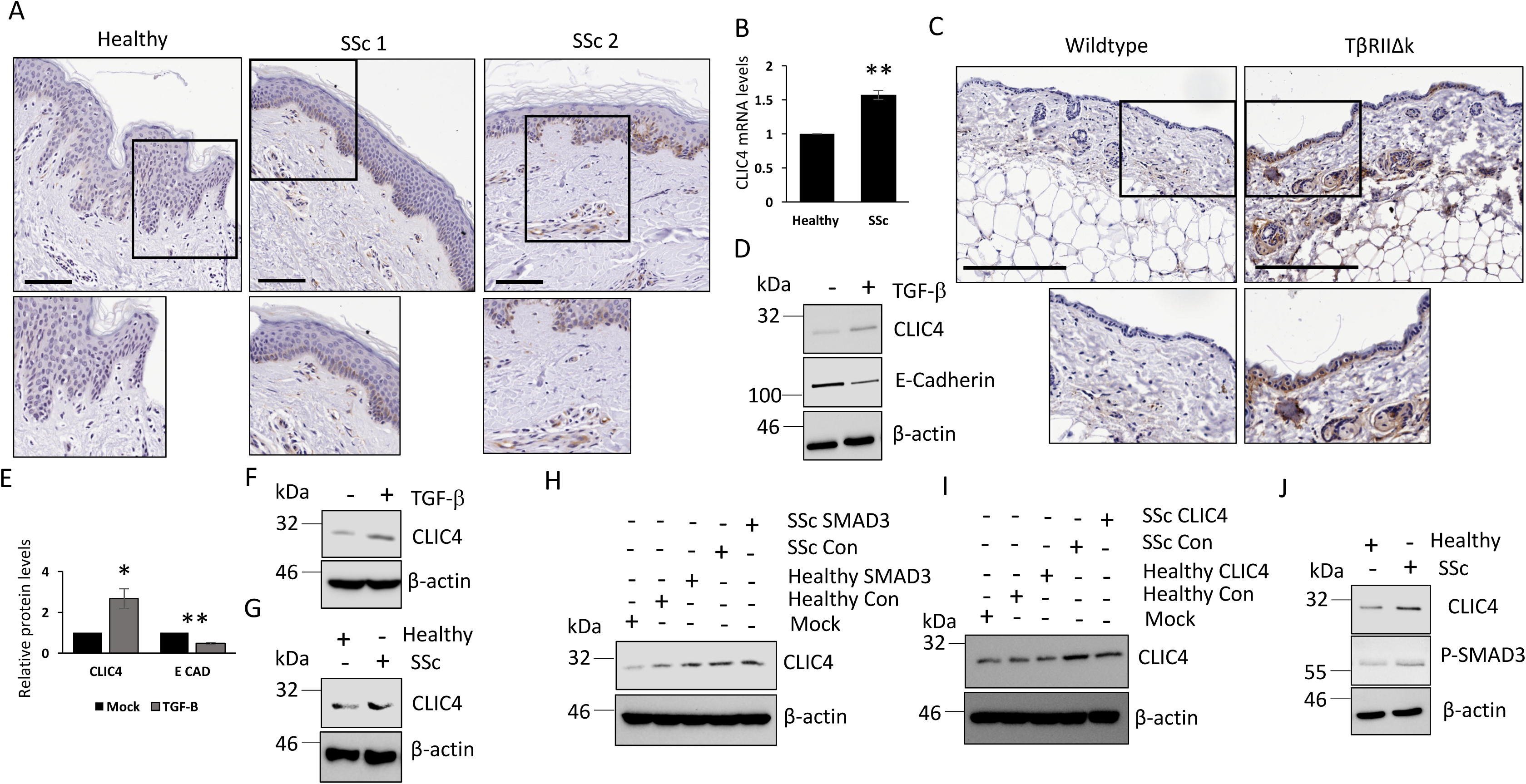
CLIC4 is elevated in SSc skin fibroblasts, keratinocytes and endothelial cells through TGF-β. (A) Skin biopsies from healthy and SSc patient forearms were stained with an antibody specific to CLIC4 and visualized with an HRP conjugated secondary antibody. The black boxes represent enlarged areas of interest in the skin. Scale bars represent 50μM. RNA was extracted from isolated SSc skin keratinocytes. (B) CLIC4 transcript levels were analysed by qPCR. (C) Immunohistochemistry analysis for CLIC4 expression (Brown) in skin sections from age matched wildtype and TβRIIΔK-fib transgenic mice. Sections were counterstained with haematoxylin (Blue). Scale bars represent 200μM. Protein was extracted from Hacats cells stimulated with TGF-β for 48 hours. (D) CLIC4 and E-cadherin protein levels were assessed by western blot. β-actin was used as a loading control. (E) Densitometry analysis of the CLIC4 and E-cadherin western blots. Protein was extracted from HUVECs endothelial cells stimulated with TGF-β for 48 hours. (F) CLIC4 protein levels were assessed by western blot. β - actin was used as a loading control. (G) Protein was extracted from Hacat cells that were grown in trans-wells in the presence of healthy or SSc dermal fibroblasts. CLIC4 protein levels were assessed by western blot. Conditioned media was collected from healthy and SSc fibroblasts transfected with scramble and SMAD3 siRNA (H) or CLIC4 siRNA (I). The media was used to stimulate Hacats. CLIC4 and β-actin protein levels were assessed by western blot. (J) HUVECs cells were stimulated with healthy and SSc fibroblast conditioned media for 24 hours. CLIC4 and pSMAD2/3 protein levels were assessed by western blot.

### TGF-β regulates CLIC4 levels in dermal keratinocytes and endothelial cells

CLIC4 transcription is regulated by the TGF-β mediated transcription factors SMAD2/3 in a range of cell types, including fibroblasts (3). To determine if TGF-β drives CLIC4 expression in skin keratinocytes and endothelial cells, we analysed by immunohistochemistry skin sections from transgenic mice with fibroblast directed activation of TGF-β signalling. As previously described the TβRIIΔk mice display all the hallmarks of skin fibrosis with enhanced TGF-β signaling characterized by increased SMAD phosphorylation in fibroblasts leading paracrine activation of other cell types. This mouse strain provides a comprehensive phenocopy of human systemic sclerosis (17). Interestingly CLIC4 levels were elevated in the fibroblasts, keratinocytes and endothelial cells of the skin in these mice (Figure1C). This suggests TGF-β signaling regulates CLIC4 expression in dermal keratinocytes and endothelial cells. To validate these finding we stimulated the human keratinocyte cell line Hacats with TGF-β and assessed CLIC4 levels. TGF-β stimulation resulted in increased CLIC4 protein (Figure1D-E). TGF-β stimulation resulted in reduced E-cadherin indicating epithelial to mesenchymal transition (EMT) was occurring, which confirms activation of the TGF-β signaling pathway. In addition we stimulated the human endothelial cell line HUVEC with TGF-β and found increased CLIC4 protein levels (Figure1F).

SSc fibroblasts are known to secrete factors that induce TGF-β signalling in neighboring cells. Therefore SSc fibroblasts may induce CLIC4 expression in neighboring keratinocytes and endothelial cells in SSc patient skin through paracrine signaling. To assess this hypothesis, Healthy and SSc dermal fibroblasts were grown in a trans-well co-culture system with Hacats. Hacats grown in culture with SSc dermal fibroblasts expressed higher levels of CLIC4 compared to Hacats grown with healthy fibroblasts (Figure1G). This suggest secreted factors from SSc fibroblasts drive CLIC4 expression in skin keratinocytes. To determine if the TGF-β signalling cascade in the fibroblasts was important for paracrine induction of CLIC4 in the keratinocytes we collected conditioned media from healthy and SSc fibroblasts that had SMAD3 depleted from the fibroblasts through siRNA knockdown. The SSc fibroblast media induced CLIC4 expression compared to mock and healthy fibroblast media (Figure1H) but depletion of SMAD3 in the SSc fibroblasts did not affect the induction of CLIC4 in the keratinocytes. The efficiency of the siRNA was confirmed through probing the fibroblast lysates for total SMAD3 (Supplementary Figure1B). This data suggests that the secretion of fibronectin and CTGF from the SSc fibroblasts does not play a role in triggering CLIC4 expression in the keratinocytes as the expression and secretion of both would be suppressed when SMAD3 was depleted. Also CLIC4 levels in the fibroblasts does not affect the ability of supernatant to induce CLIC4 expression in the keratinocytes as SMAD3 knockdown reduced CLIC4 expression levels in the SSc fibroblasts (Supplementary Figure1B). This was confirmed when we observed no change in CLIC4 expression levels in Hacats stimulated with conditioned media from SSc fibroblasts transfected with scramble of CLIC4 siRNA (Figure1I). CLIC4 protein levels were elevated in Hacats stimulated with SSc fibroblasts media compared to healthy control fibroblast media and knockdown of CLIC4 in the SSc fibroblast did not affect the ability of the SSc fibroblast media to induce CLIC4 expression in the Hacats. We confirmed efficient CLIC4 knockdown in the healthy and SSc fibroblasts by western blot (Supplementary Figure2A). Interestingly the knockdown of CLIC4 was also able to disrupt the pro-fibrotic phenotypes in SSc fibroblasts. CLIC4 knockdown blocked pro-fibrotic gene expression in SSc dermal fibroblasts (Supplementary Figure2). The CLIC4 siRNA reduced CLIC4 protein (Supplementary Figure2A-B) and transcript levels (Supplementary Figure2C) in both healthy and SSc fibroblasts. The disruption in CLIC4 expression resulted in reduction of the pro-fibrotic signalling factors β-catenin (Supplementary Figure2A, B and D), pSMAD3 (Supplementary Figure2A-B) and GLI2 (Supplementary Figure2A, E) as well as expression of the pro-fibrotic markers CTGF and α-SMA in SSc fibroblasts but had no effect in healthy fibroblasts. These results validated previous data from our group where the CLIC4 inhibitors NPPB and IAA-94 block pro-fibrotic gene expression as well as the Wnt3a and hedgehog signaling pathways in SSc dermal fibroblasts (3,4).

**Figure 2:**
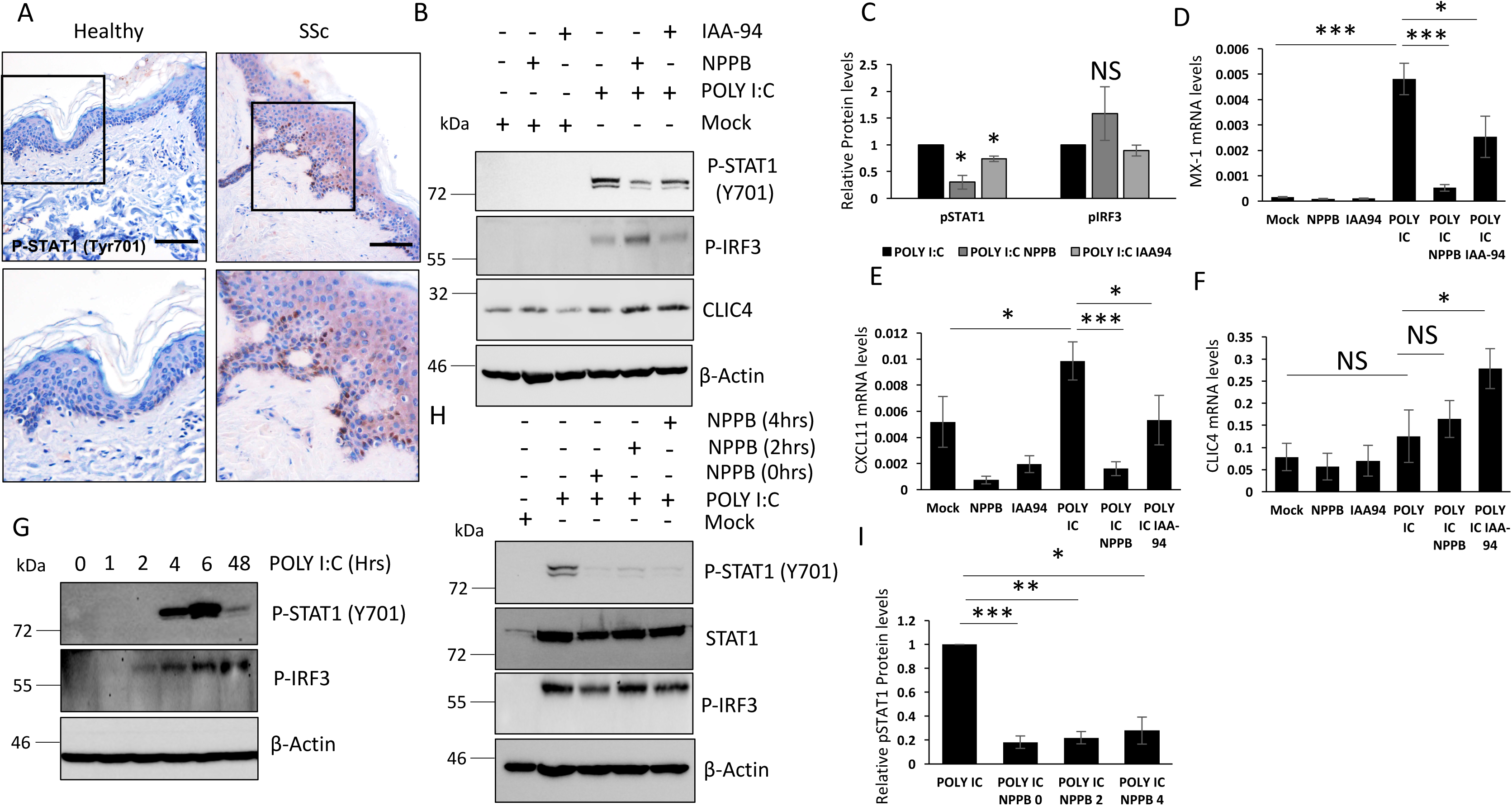
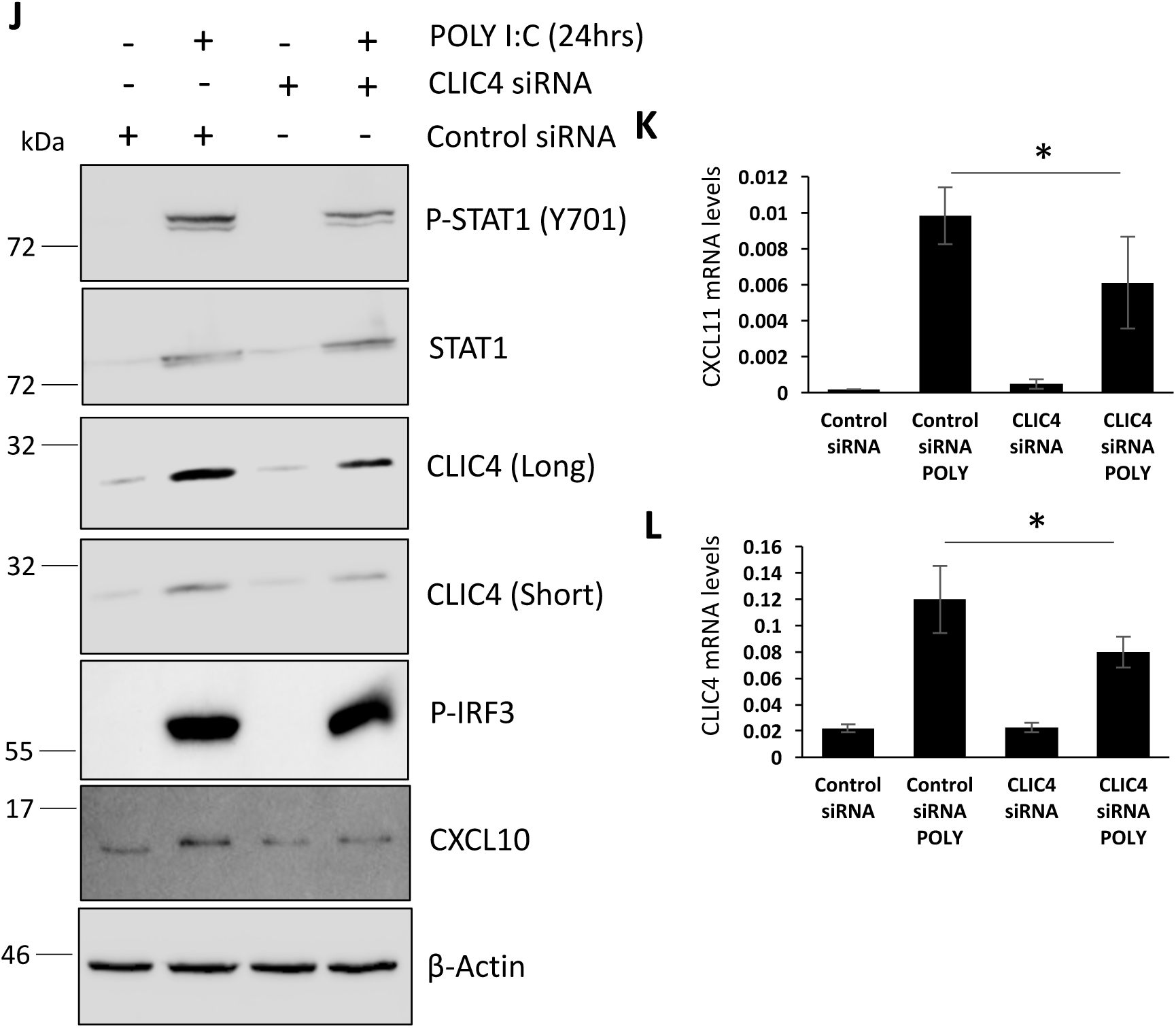
CLIC4 plays an important role in TLR3 mediated STAT1 activation in keratinocytes. (A) Skin biopsies from healthy and SSc patient forearms were stained with an antibody specific to phosphorylated STAT1 and visualized with an HRP conjugated secondary antibody. The black boxes represent enlarged areas of interest in the skin. Protein and RNA were extracted from Hacats stimulated with POLY I:C for 48 hours. In addition, Hacats were treated with the chloride channel inhibitors NPPB and IAA-94. (B) pSTAT1, pIRF3 and CLIC4 protein levels were assessed by western blot. β -actin was used as a loading control. (C) Graphs represent densitometry analysis for the mean and standard error for three independent experiments. MX-1 (D), CXCL11 (E) and CLIC4 (F) transcript levels were assessed by qPCR. Protein was extracted from Hacats stimulated with POLY I:C for 1, 2, 4, 6 and 48 hours. (G) pSTAT1 and pIRF3 protein levels were assessed by western blot. β -actin was used as a loading control. Protein was extracted from Hacats stimulated with POLY I:C for 48 hours. In addition the Hacat cells were treated with NPPB at the same time poly I:C and 2, 4 hours post POLY I:C stimulation. (H) pSTAT1, total STAT1 and pIRF3 protein levels were assessed by western blot. β-actin was used as a loading control. (I) Graphs represent densitometry analysis for the mean and standard error for three independent experiments. Protein and RNA were isolated from Hacats cells transfected with siRNA specific for CLIC4 or a scramble control siRNA for 48 hours. 24 hours prior to harvesting the cells were stimulated with POLY I:C. (J) pSTAT1, Total STAT1, pIRF3, CXCL10 and CLIC4 proteins levels were assessed by western blot. β -actin was used as a loading control. CXCL11 (K) and CLIC4 (L) transcript levels were assessed by qPCR.

We also observed increased CLIC4 expression in HUVEC endothelial cells grown in SSc fibroblast conditioned media compared to cells grown in healthy fibroblast conditioned media (Figure1J). In addition pSMAD3 levels were increased in the HUVECs grown in SSc fibroblast media confirming activation of the TGF-β signaling pathway. This suggests SSc fibroblasts may play a central role in regulating TGF-β signaling and in turn CLIC4 expression in neighboring cells within the skin.

SSc keratinocytes are characterized by aberrant EMT which is regulated in part by the TGF-β signalling pathway (18). As we have shown CLIC4 and TGF-β are inherently linked in keratinocytes it is possible CLIC4 may induce EMT in the SSc keratinocytes. To test this hypothesis we stimulated Hacats with TGF-β in the presence and absence of the chloride channel inhibitors NPPB and IAA-94. Both these inhibitors have previously been shown to block CLIC4 activity (3, 4). The TGF-β induced EMT as shown through the reduction of E cadherin expression. The chloride channel inhibitors did not affect the TGF-β mediated reduction in E-Cadherin (Supplementary Figure3A). Similar results were obtained with CLIC4 siRNA. Hacats were transfected with siRNA specific to CLIC4 and then stimulated with TGF-β. The TGF-β stimulation reduced E-Cadherin levels to similar levels in both the scramble siRNA and CLIC4 siRNA cells (Supplementary Figure3B). This suggests CLIC4 does not have a role in the aberrant EMT observed in SSc keratinocytes.

**Figure 3:**
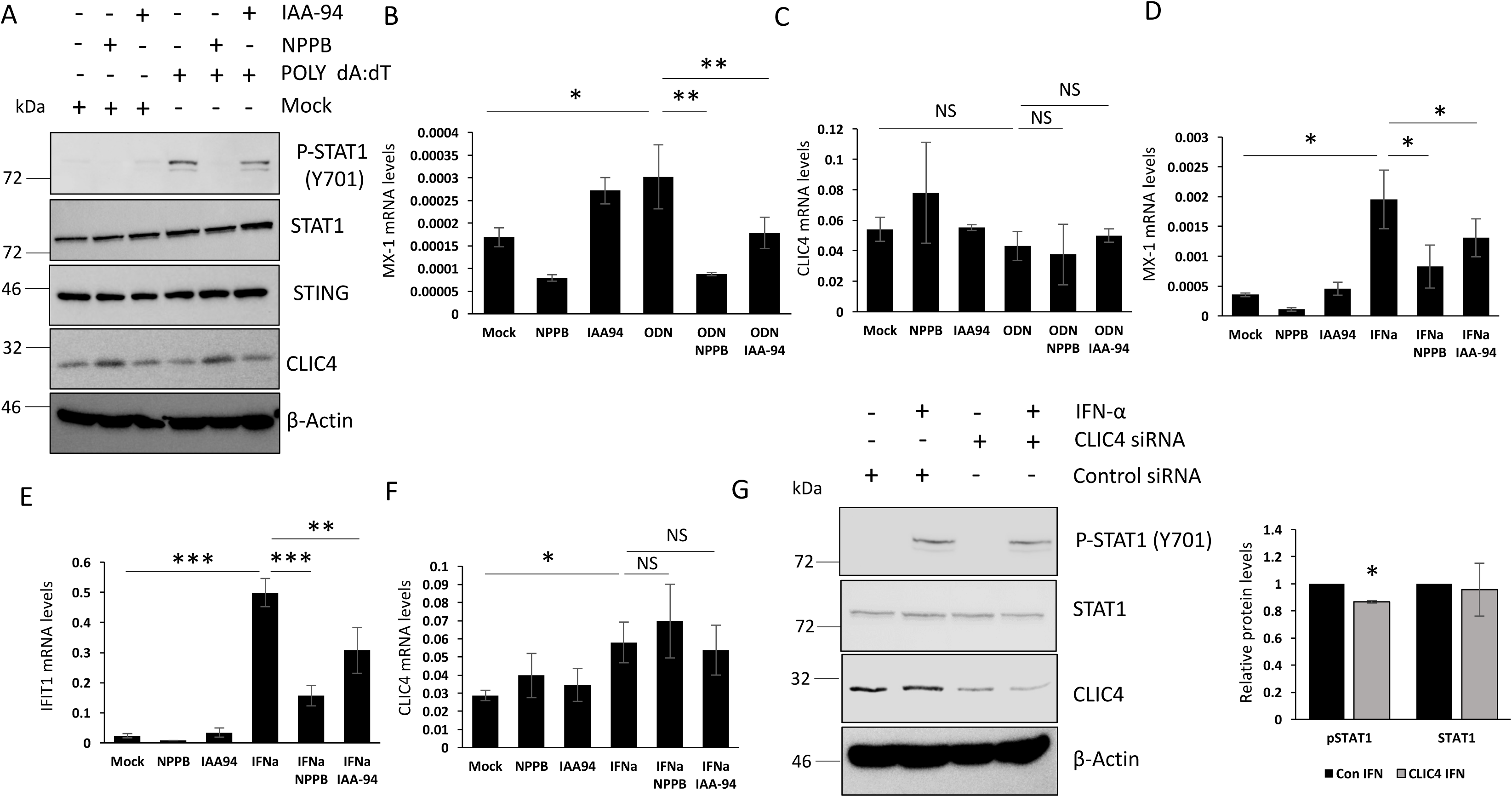
CLIC4 plays an important role for the activation of the Type 1 interferon pathway in keratinocytes. (A) Protein was extracted from Hacats stimulated with POLY dA:dT for 48 hours. In addition, Hacats were treated with the chloride channel inhibitors NPPB and IAA-94. pSTAT1, Total STAT1, TOTAL STING and CLIC4 protein levels were assessed by western blot. β-actin was used as a loading control. RNA was extracted from Hacats stimulated with ODN2216 for 48 hours. In addition, Hacats were treated with the chloride channel inhibitors NPPB and IAA-94. MX-1 (B) and CLIC4 (C) transcript levels were assessed by qPCR. RNA was extracted from Hacats cells treated with the chloride channel inhibitors NPPB and IAA94 for 48 hours. In addition the Hacats were stimulated with IFN2-α for 2.5 hours prior to harvesting. MX-1 (D), IFIT1 (E) and CLIC4 (F) transcript levels were assessed by qPCR. (G) Protein was extracted from Hacats transfected with scramble and CLIC4 siRNA. In addition the cells were stimulated with IFN-2alpha for 2.5hours. pSTAT1, Total STAT1 and CLIC4 protein levels were assessed by western blot. β -actin was used as a loading control. Graph represent pSTAT1 densitometry analysis for the mean and standard error for three independent experiments

### Inhibition of CLIC4 plays an important role in regulating STAT1 mediated ISG expression in keratinocytes

Keratinocytes in SSc skin are a major driver of the type-1 IFN signaling in SSc skin (6) and display high levels of nuclear phosphorylated STAT1 compared to healthy control skin (Figure2A). STAT1 is a major transcription factor associated with the type-1 interferon signaling pathway. Previous studies have shown that CLIC4 is important in triggering inflammation. CLIC4 along with CLIC1 can trigger the activation of the NLRP3 inflammasome and IL-1β in macrophages (19, 20), suggesting that CLIC4 may be directly involved in the molecular events sustaining inflammation. Therefore, we investigated if CLIC4 could play a role in the type-1 IFN signalling observed in SSc keratinocytes. Hacat cells were stimulated with a number of immune stimuli that trigger a type-1 IFN response in the presence and absence of the chloride channel inhibitors NPPB and IAA-94. The TLR3 agonist POLY I:C (48 hours) was able to induce STAT1 activation (Phosphorylation Y701) in Hacats and interestingly both chloride channel inhibitors attenuated STAT1 activation suggesting CLIC4 plays a role in the type-1 IFN response in keratinocytes (Figure2B-C). Next, we investigated if the inhibition of STAT1 resulted in downstream inhibition of Interferon stimulated genes (ISGs). Blocking the activity of CLIC4 resulted in reduced MX1 and CXCL11 expression in POLY I:C stimulated cells (Figure2D-E). This confirms NPPB and IAA-94 ability to block STAT1 activation resulted in a reduction in downstream signaling. TLR3 activation did not induce expression of CLIC4 in keratinocytes. CLIC4 protein (Figure2B) and transcript (Figure2F) levels were not altered in keratinocytes stimulated with POLY I:C for 48 hours.

The Interferon regulatory transcription factor 3 (IRF3) is known to be an important component of the type-1 interferon signaling cascade and is found upstream of STAT1 in the cascade. The chloride channel inhibitors did not inhibit IRF3 phosphorylation after POLY I:C stimulation (Figure2B-C). This suggests CLIC4 regulates STAT1 activation downstream of IRF3. To further assess where CLIC4 intervenes in the cascade, we inhibited CLIC4 after the activation of IRF3 and STAT1. Since IRF3 and STAT1 were activated 2- and 4-hours post POLY I:C stimulation respectfully (Figure2G), we inhibited CLIC4 with NPPB, 2 and 4 hours post POLY I:C stimulation (Figure2H). In cooperation with the data presented in Figure 2B, NPPB blocked STAT1 activation after the activation of IRF3 (2hrs). Interestingly NPPB reduced the phosphorylated STAT1 levels after its initial activation (4hrs) (Figure2H-I). This data rules out a role for CLIC4 in regulating IFN expression or secretion and Interferon receptor activation and suggests inhibition of CLIC4 disrupts the STAT1 phosphorylation.

### Depletion of CLIC4 plays an important role in regulating STAT1 mediated ISG expression in keratinocytes and pro-fibrotic gene expression in fibroblasts

The chloride channel inhibitors NPPB and IAA-94 target a number of other chloride channels in addition to CLIC4. Therefore to rule out off-target effects of the inhibitors we repeated the experiments above with siRNA specific to CLIC4. Hacat cells were transfected with CLIC4 siRNA for 24 hours and then the cells were stimulated with POLY I:C for a further 24 hours. We observed a 45-50% knockdown in CLIC4 protein and transcript levels (Figure2J, L) in cells transfected with the CLIC4 siRNA. This knockdown in CLIC4 resulted in an attenuation of STAT1 activation by POLY I:C (30% reduction) (Figure2J). This reduction in STAT1 activation lead to a reduction in ISG gene expression (CXCL11 (35% reduction)) (Figure2K). In keeping with the inhibitor data knockdown of CLIC4 did not affect IRF3 activation in POLY I:C stimulated Hacats (Figure2J). Interestingly we observed an increase in CLIC4 protein expression levels (Figure2J) when the Hacats were stimulated with POLY I:C for 24 hours whereas we observed no change in CLIC4 levels after 48 hours stimulation (Figure2B). As CLIC4 protein levels change depending on the time of addition of POLY IC, we assessed CLIC4 protein levels across the POLY I:C time course (Supplementary Figure4). Interestingly we found no change in CLIC4 expression levels in Hacats stimulated with POLY I:C for a short period of time (1-6hours). We observed increased CLIC4 protein levels at 24 hours and levels returned to unstimulated levels at 48 hours. This suggests CLIC4 protein levels transiently changes during the course of the stimulation.

**Figure 4:**
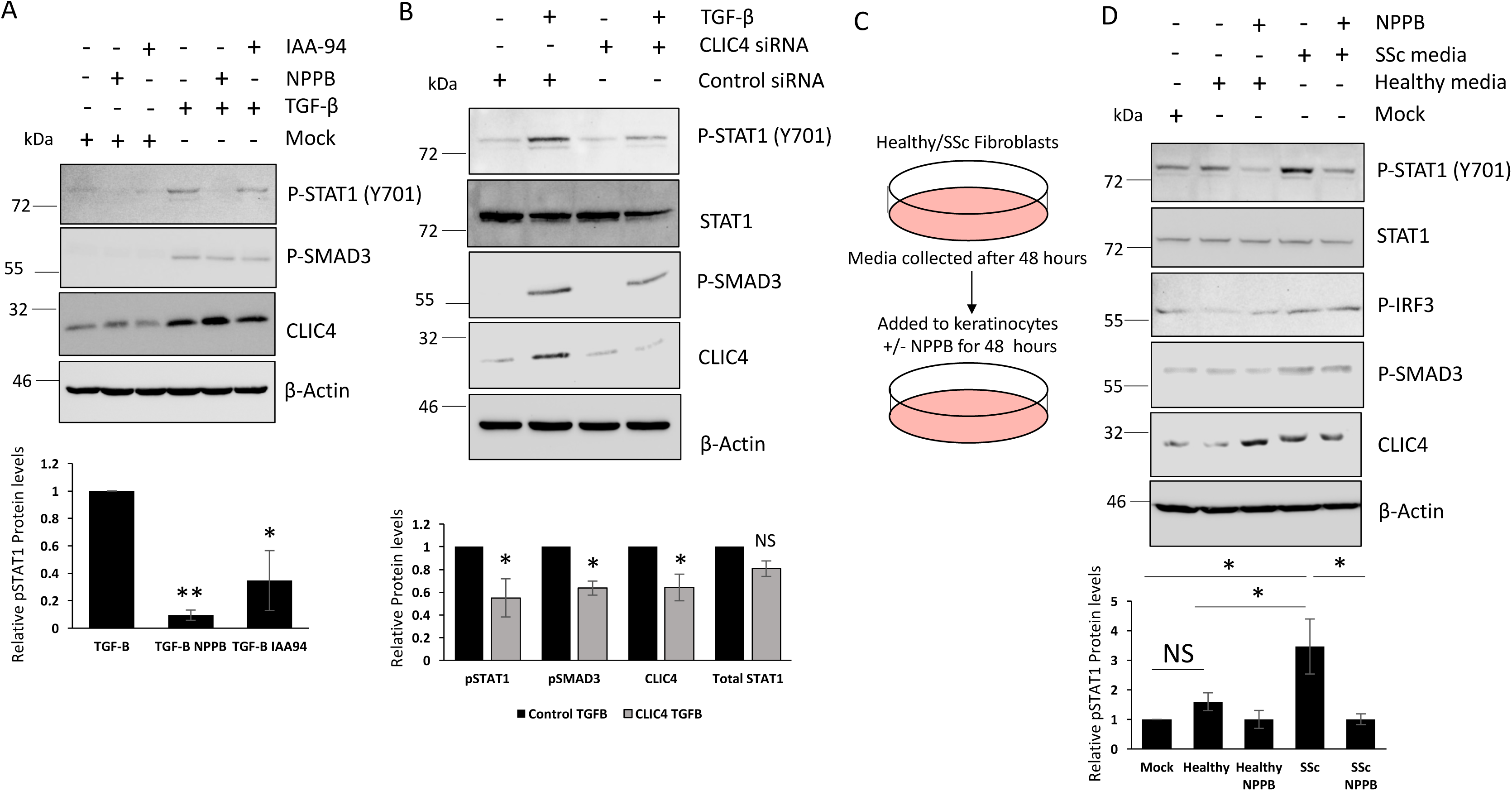
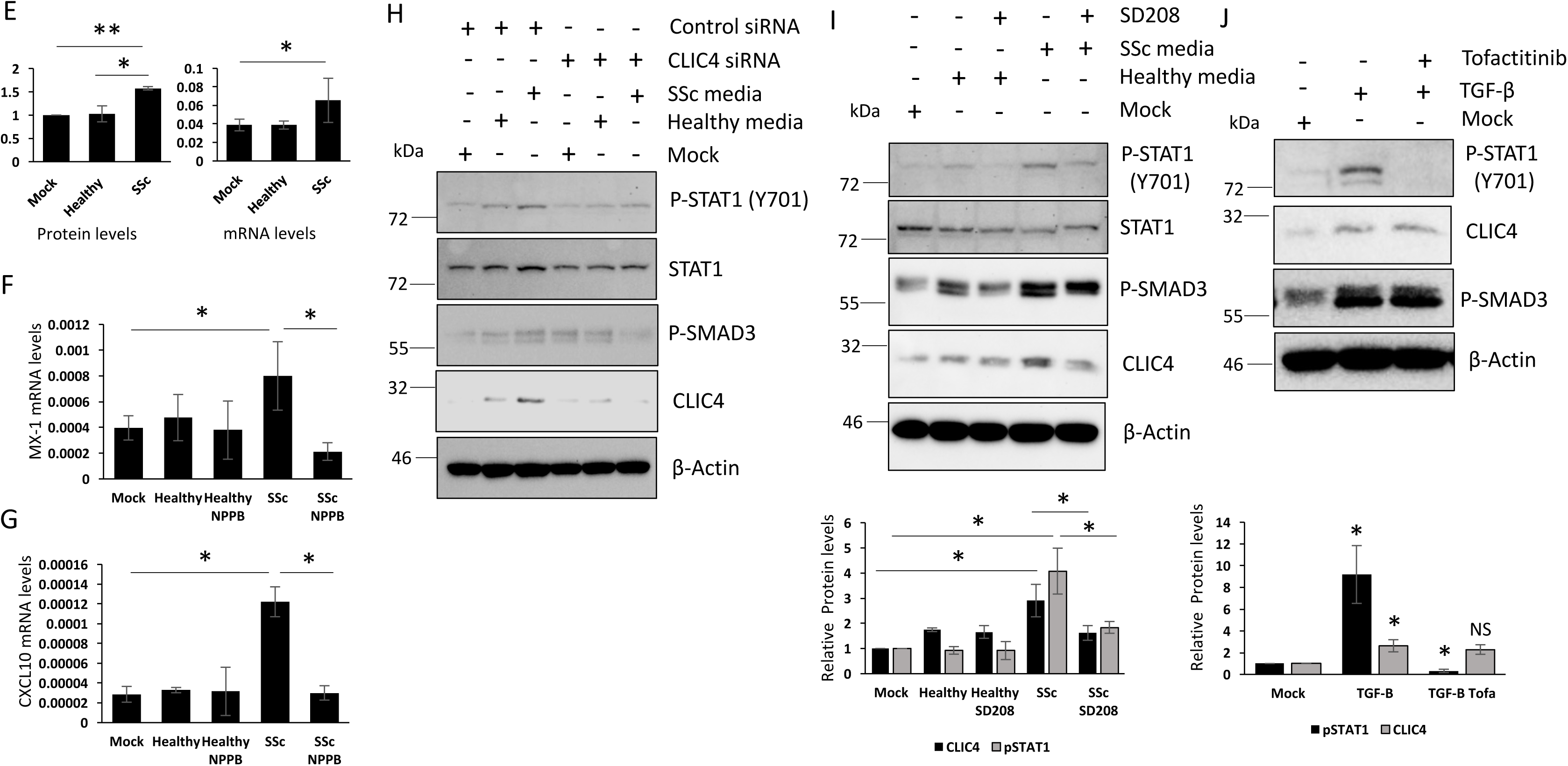
Inhibition of CLIC4 blocks SSc derived STAT1 activators. Protein was extracted from Hacats stimulated with TGF-β for 48 hours. In addition, Hacats were treated with the chloride channel inhibitors NPPB and IAA-94. (A) pSTAT1, CLIC4 and pSMAD2/3 protein levels were assessed by western blot. β-actin was used as a loading control. Graphs represent densitometry analysis for the mean and standard error for three independent experiments. (B) Protein was extracted from, Hacats were transfected with the scramble and CLIC4 siRNA. After 24 hours the transfected Hacats were stimulated with TGF-β for a further 24 hours. pSTAT1, total STAT1, pSMAD2/3 and CLIC4 protein levels were assessed by western blot. β-actin was used as a loading control. Graphs represent densitometry analysis for the mean and standard error for three independent experiments. (C) Serum depleted conditioned media was collected from healthy and SSc patient fibroblasts after 48hrs. Hacats were stimulated with the media for 48 hours in the absence or presence of NPPB. (D) pSTAT1, STAT1 pIRF3, pSMAD3 and CLIC4 protein levels were assessed by western blot. β-actin was used as a loading control. Graph represent densitometry analysis for the mean and standard error for three independent experiments. (E) CLIC4 protein and mRNA levels were assessed in Hacats stimulated with healthy and SSc fibroblast conditioned media. MX1 (F) and CXCL10 (G) transcript levels were assessed by qPCR in Hacats stimulated with healthy and SSc fibroblast conditioned media plus NPPB. (H) Hacats were transfected with scramble or CLIC4 siRNA for 24 hours. Then stimulated with healthy or SSc fibroblast conditioned media for a further 48 hours. pSTAT1, STAT1, pSMAD2/3 and CLIC4 protein levels were analysed by western blot. β -actin was used as a loading control. Hacats were stimulated with healthy and SSc fibroblast media for 48 hours in the absence or presence of the TGF-β receptor inhibitor SD208. (I) pSTAT1, total STAT1 pSMAD2/3 and CLIC4 protein levels were assessed by western blot. β-actin was used as a loading control. Graph represent densitometry analysis for the mean and standard error for three independent experiments. Hacats were stimulated with TGF-β in the absence or presence of the JAK1 inhibitor Tofacanib for 48 hours. (J) pSTAT1, pSMAD2/3 and CLIC4 protein levels were assessed by western blot. β-actin was used as a loading control. Graph represent densitometry analysis for the mean and standard error for three independent experiments.

### CLIC4 regulates type-1 IFN signaling induced by TLR9 and cytosolic DNA sensors in keratinocytes

Next we investigated whether the inhibition of CLIC4 prevented STAT1 activation and ISG gene expression in the context of other immune stimuli or if this was specific for TLR3 (POLY I:C) signaling. We observed similar results when Hacat cells were stimulated with the cytosolic DNA sensor agonist POLY dA:dT. Poly dA:dT stimulation resulted in activation of STAT1 in Hacats (Figure3A). This was blocked in the presence of both chloride channel inhibitors. Poly dA:dT stimulation did not affect CLIC4 expression levels in Hacats. STING is a major downstream regulator of the cytosolic DNA sensor complexes. Interestingly the chloride channel inhibitors did not affect the expression levels of STING in Hacat cells. In collaboration with the POLY I:C data this suggests that CLIC4 specifically regulates STAT1 phosphorylation and does not affect upstream factors regardless of the stimuli. Inhibition of CLIC4 blocked TLR9 mediated ISG expression. We observed increased expression of MX1 in ODN2216 (TLR9 agonist) treated cells, which was blocked by the chloride channel inhibitors (Figure3B). CLIC4 gene expression levels were not altered in ODN2216 stimulated Hacats (Figure3C). Inhibition of CLIC4 blocked the induction of MX1 and IFIT1 by IFN-2alpha in Hacats. Hacats were pre-treated with NPPB and IAA-94 for 45.5 hours and the cells were stimulated with IFN-2alpha for 2.5 hours prior to harvesting. IFN-2alpha increased MX1 and IFIT1 expression and this was partially blocked with both inhibitors (Figure3D-E). Interestingly stimulation of Hacats with IFN-alpha for 2.5hrs resulted in increased CLIC4 gene expression (Figure3F), further highlighting the transient nature of CLIC4 expression upon stimulation with type-1 IFN agonists. Depletion of CLIC4 from Hacats had similar effects to the inhibitors. Depletion of CLIC4 in Hacats attenuated the activation of STAT1 (15%) in response to IFN-2alpha (2.5 hour stimulaton) compared to scramble control cells stimulated with IFN-2alpha (Figure3G). Taken together the data here shows CLIC4 plays an important role in regulating the type-1 interferon signaling cascade regardless of the stimuli.

### Inhibition of CLIC4 blocks SSc derived STAT1 activators

The data described above shows inhibition of CLIC4 blocked STAT1 activation by a range of immune stimuli. We next wanted to investigate the ability of the CLIC4 inhibitors to block STAT1 activation by SSc derived triggers such as TGF-β or SSc fibroblast supernatant. TGF-β is a major cytokine involved in the pathogenesis associated with SSc. TGF-β was able to induce STAT1 activation in Hacats and this was reversed when CLIC4 is inhibited with the chloride channel inhibitors (Figure4A) or the CLIC4 siRNA (Figure4B).

Previous studies have shown SSc fibroblasts can trigger the activation of STAT1 in keratinocytes (6). We therefore investigated if CLIC4 was involved in this process. Conditioned media from healthy and SSc dermal fibroblasts was used to stimulate Hacat cells for 48 hours plus/minus the chloride channel inhibitor NPPB (Figure4C). We observed increased STAT1 phosphorylation in keratinocytes stimulated with SSc fibroblast media compared to mock and healthy fibroblast media (Figure4D). This was reversed when the keratinocytes were treated with the CLIC4 inhibitor NPPB. The inhibition of STAT1 lead to a 75% reduction in MX1 and CXCL10 transcript levels in Hacats stimulated with SSc fibroblast media plus NPPB (Figure4F-G). In addition we observed an increase in phosphorylated IRF3 levels in the keratinocytes stimulated with the SSc fibroblast media but this was not altered with the addition of NPPB (Figure4D), further confirming that CLIC4 does not regulate IRF3 activation. We also observed that the SSc fibroblast media induced a 1.5-fold increase in CLIC4 protein (Figure4D) and transcript levels (Figure4E) in the keratinocytes. This further validates the trans-well co-culture experiments in Figure1. Similar results were observed when conditioned media from primary healthy and SSc fibroblasts was used. Primary SSc fibroblast media induced CLIC4 expression as well as STAT1 and SMAD2/3 activation in HaCats. The activation was reversed with NPPB (Supplementary Figure5A). Depletion of CLIC4 in HaCats using siRNA blocked the ability of the SSc fibroblast supernatant to induce the phosphorylation STAT1 and SMAD2/3. (Figure4H). Depletion of CLIC4 in the SSc fibroblasts did not attenuate the ability of the SSc fibroblast media to induce the activation of CLIC4 (Supplementary Figure5B). This is unsurprising as the depletion of CLIC4 in the SSc fibroblasts was previously shown to not affect the ability of the fibroblast media to induce CLIC4 expression in the Hacats (Figure1I).

**Figure 5:**
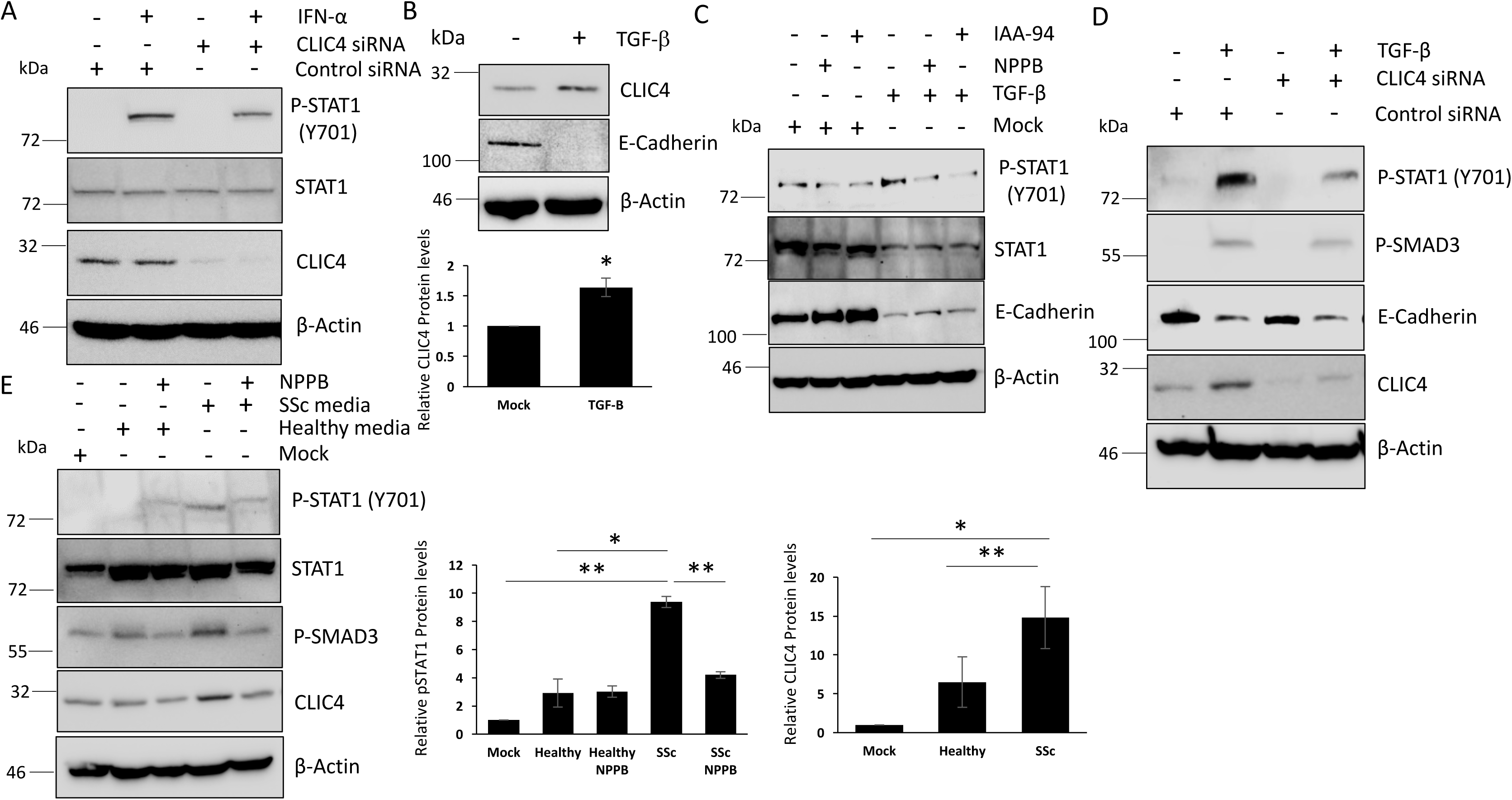
Inhibition of CLIC4 prevents STAT1 activation in lung epithelial cells. Protein was extracted from A549 cells transfected with CLIC4 siRNA for 72 hours. In addition the A549 cells were stimulated with IFN2-α for 2.5 hours prior to harvesting. pSTAT1, STAT1 and CLIC4 protein levels were assessed by western blot. β -actin was used as a loading control. Protein was extracted from A549 stimulated with TGF-β for 48 hours. (B) CLIC4 and E-Cadherin protein levels were assessed by western blot. β -actin was used as a loading control. Graph represent densitometry analysis for the mean and standard error for three independent experiments. Protein was extracted from A549 stimulated with TGF-β for 48 hours. In addition, A549 cells were treated with the chloride channel inhibitors NPPB and IAA-94. (C) pSTAT1, total STAT1 and E-Cadherin protein levels were assessed by western blot. β -actin was used as a loading control. Protein was extracted from A549 transfected with CLIC4 siRNA for 72 hours. In addition, the A549 cells were stimulated with TGF-β for the final 24 hours. (D) pSTAT1, pSMAD2/3, CLIC4 and E-cadherin protein levels were assessed by western blot. β -actin was used as a loading control. (E) Serum depleted conditioned media was collected from healthy and SSc patient fibroblasts after 48hrs. A549 cells were stimulated with the media for 48 hours in the absence or presence of NPPB. pSTAT1, STAT1, pSMAD3 and CLIC4 protein levels were assessed by western blot. β-actin was used as a loading control. Graphs represent CLIC4 and pSTAT1 densitometry analysis for the mean and standard error for three independent experiments

SSc fibroblast media had the ability to induce STAT1 activation and CLIC4 expression in the keratinocytes. We therefore explored the mechanism behind the induction of CLIC4. SSc fibroblast exosomes have previously been shown to play an important role in the regulating the type 1 interferon response in the keratinocytes (6). Therefore we assessed if they played a role in regulating CLIC4 expression. CLIC4 expression levels were unaffected in keratinocytes stimulated with healthy and SSc fibroblast exosomes (Supplementary Figure6). This suggests additional factors within the SSc fibroblast supernatant drive CLIC4 expression in the keratinocytes and CLIC4 induces a type-1 IFN response in keratinocytes independent of the exosomes.

**Figure 6:**
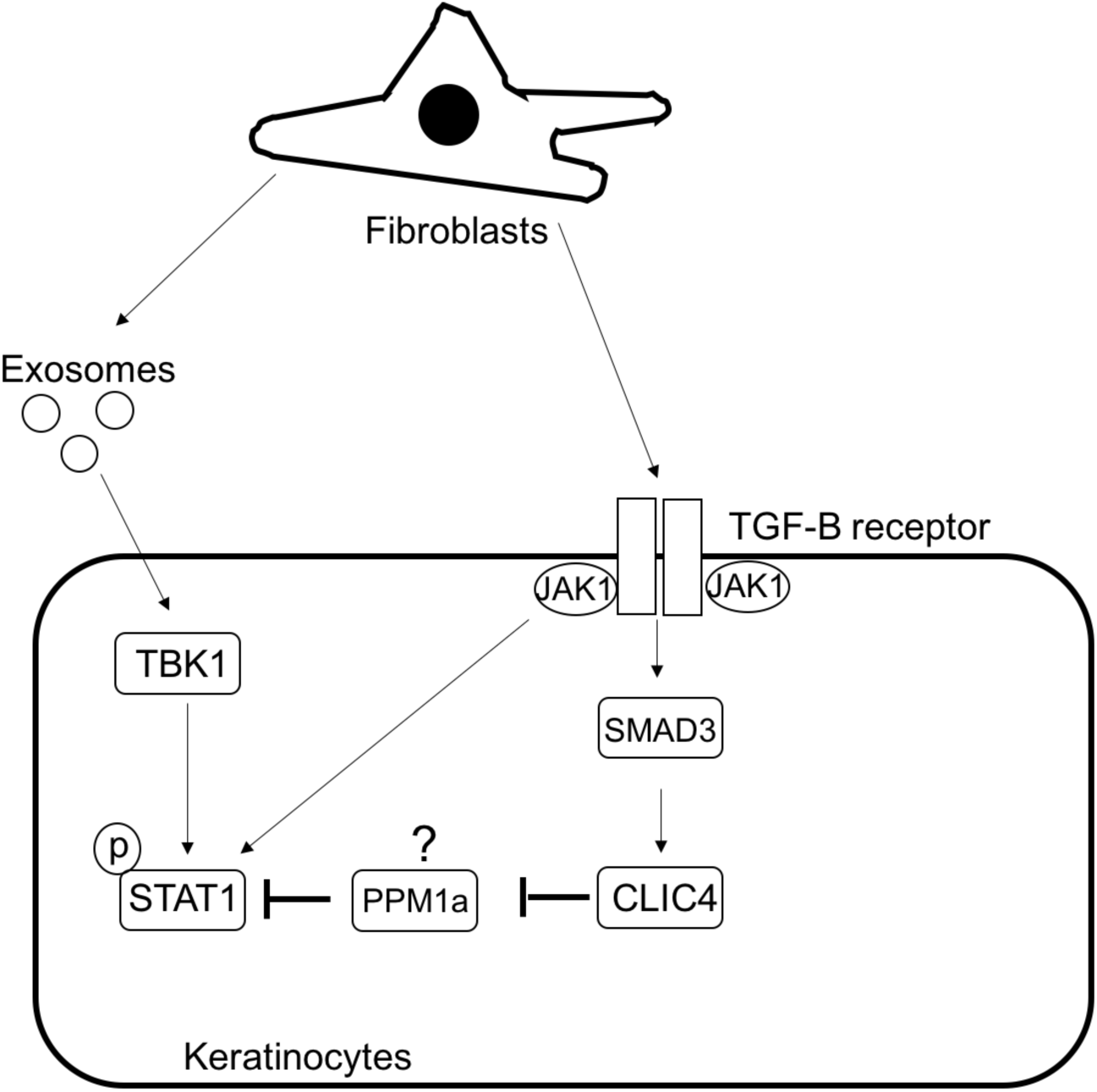
CLIC4 regulates STAT1 activation in epithelial cells through a TGF-β dependent pathway.

As described above TGF-β can induce STAT1 activation in the Hacats (Figure4A-B) and SSc fibroblast media is able to activate SMAD2/3 in the Hacats (Figure4D, H) and the activation of both is attenuated when CLIC4 is inhibited in the Hacats. This suggests that SSc fibroblast supernatant may induce STAT1 activation through the activation of the TGF-β receptor in Hacats. To test this hypothesis we stimulated keratinocytes with healthy and SSc fibroblast media with the addition of the TGF-β receptor inhibitor SD208 (Figure4I). The SSc fibroblast media increased CLIC4 and pSMAD3 levels in the keratinocytes and this was reversed in the presence of SD208. Interestingly SD208 blocked the induced of pSTAT1 in keratinocytes stimulated with SSc fibroblast media (Figure4I). Further analysis revealed TGF-β receptor mediated activation of STAT1 was JAK dependent. We were able to reverse the activation of STAT1 in Hacats stimulated with TGF-β in the presence of the specific JAK inhibitor Tofacitinib (Figure4J). Interestingly Tofacitinib did not block TGF-β mediated activation of SMAD2/3 or increased expression of CLIC4 (Figure4J). This suggests activation of the TGF-β receptor by SSc fibroblast media induces CLIC4 expression in the keratinocytes through SMAD2/3 and STAT1 is activated by SSc fibroblast media through interplay between the TGF-β receptor and JAK.

### Disrupting CLIC4 activity blocks Type 1 Interferon signaling in Lung epithelial cells

Previous studies have shown the aberrant type-1 IFN signalling is a contributing factor to SSc-Interstitial lung disease (SSc-ILD) (21). The lung epithelial cells play a major role in the type-1 IFN response induced by a number of viruses and injury. The lung epithelial cells have been shown to be a driver of SSc-ILD (22). Therefore, we investigated the role CLIC4 plays in type 1 IFN signaling in the lung epithelial cell line A549. CLIC4 was required for IFN-alpha mediated STAT1 activation in A549 cells. We observed an induction of p-STAT1 in control siRNA cells stimulated with IFN-alpha and this was attenuated with the CLIC4 siRNA (Figure5A). Similar to Hacats, CLIC4 expression was regulated by TGF-β in the lung epithelial cell line A549. A549 cells stimulated with TGF-β induced the expression of CLIC4 (1.6-fold) (Figure5B) and inhibition of CLIC4 with the chloride channel inhibitors NPPB and IAA-94 (Figure5C) or CLIC4 siRNA (Figure5D) blocked TGF-β mediated STAT1 activation in these cells. Similar to keratinocytes, inhibition of CLIC4 did not alter TGF-β mediated EMT in the lung epithelial cells (Figure5C-D). We observed a reduction in E cadherin levels in A549 cells stimulated with TGF-β and these levels were unaffected when CLIC4 was inhibited with the chloride channel inhibitors or siRNA. This is important as EMT is a major driving factor in SSc ILD progression (23). Further analysis revealed that SSc fibroblast supernatant induced CLIC4 (2.3-fold) expression levels as well as the activation of STAT1 (3.2-fold) and SMAD3 in A549 cells compared to healthy control media. The activation of STAT1 and SMAD3 was reversed with the chloride channel inhibitor NPPB (Figure5E). Taken together, this data suggests CLIC4 is an important regulator of STAT1 in a range of epithelial cell types (Keratinocytes and lung epithelial cells)

## Discussion

We have shown for the first time CLIC4 is upregulated in a number of cell types within SSc patient skin and this is driven by TGF-β. CLIC4 levels were elevated in the keratinocytes and endothelial cells of TβRIIΔk mice consistent with *in vivo* paracrine TGF-β dependent modulation of critical cells relevant to SSc pathogenesis, as has been previously demonstrated from pulmonary epithelial and endothelial compartments in this mouse strain (24). We identified cell to cell communication between the fibroblasts, keratinocytes and endothelial cells was important for regulating CLIC4 expression. Keratinocytes grown in a trans-well co-culture system with SSc fibroblasts induced CLIC4 expression (Figure1C) and conditioned media from SSc fibroblasts was able to induce CLIC4 expression in keratinocytes and endothelial cells. This was dependent on the activation of the TGF-β receptor in the keratinocytes as the TGF-β receptor inhibitor SD-208 blocked CLIC4 expression induced by the SSc fibroblast conditioned media (Figure4I).

We have shown that blocking CLIC4 with small molecule inhibitors and siRNA disrupts type-1 IFN signalling triggered by a number of immune stimuli in epithelial cells. The chloride channel inhibitors NPPB and IAA-94 and CLIC4 siRNA prevented TLR3 (POLY I:C), cGAS (POLY dA:dT), TLR9 (ODN2216) and IFN2-α mediated activation of STAT1 and downstream ISG expression (Figure 3,5). This data suggests CLIC4 is a potential global regulator of type-1 IFN signalling in epithelial cells.

Interestingly we believe CLIC4 modulates the pathway by enhancing STAT1 phosphorylation. The CLIC4 inhibitors blocked the phosphorylation of STAT1 but do not affect upstream proteins such as IRF3. Furthermore time course experiments (Figure2G-H) showed the addition of the inhibitors after the activation of STAT1 (4 hours) by POLY I:C lead to a blockage of said activation of STAT1. This is further evidence that CLIC4 regulates the type-1 interferon signaling through STAT1.

CLIC4 has previously been shown to regulate the responses of macrophages to microbial products (19). LPS stimulates CLIC4 expression in macrophages which in turn enhances the macrophages response to LPS. Interestingly CLIC4 participates in LPS induced activation of IRF3 but not STAT1 (19). This data further highlights the importance of CLIC4 in immune responses but shows it has differing roles between cell types, since in keratinocytes CLIC4 modulate STAT1, but not IRF3 activation.

The ability of the CLIC4 inhibitors to block SSc fibroblast mediated STAT1 activation in the keratinocytes is intriguing. SSc fibroblast conditioned media and exosomes have previously been shown to induce STAT1 activation through the TBK1 (6). Data presented in this study shows CLIC4 is not an intermediator of this pathway as CLIC4 expression levels were unaffected in keratinocytes stimulated by SSc fibroblast exosomes. Our hypothesis is the SSc fibroblast exosomes trigger the activation of STAT1 in keratinocytes and CLIC4 amplifies or maintain this activation leads to the dysregulated type-1 IFN signaling in SSc skin (Figure6).

Furthermore we have shown for the first time that SSc fibroblasts induce the type-1 IFN response in keratinocytes in part through the activation of the TGF-β signalling cascade and this activation is mediated in part through JAK. This is interesting as previous studies have shown JAK1 can bind to the TGF-β receptor and induce STAT activation in a SMAD independent manner (25). This corroborates data presented in this study where TGF-β mediated activation of STAT1 is disrupted by the JAK inhibitor Tofactitinib but the inhibitor does not affect SMAD2/3 activation or CLIC4 expression (Figure 4J). At the same time the SSc fibroblast media induces CLIC4 expression through TGF-β/SMAD dependent signaling. CLIC4 could then enhance STAT1 activation through the inhibition of the protein phosphatase 1a (PPM1a) (Figure6). Nuclear CLIC4 binds to PPM1a and prevents de-phosphorylation of p38 in fibroblasts (26) thus enhancing TGF-β signaling in both cell types. Interestingly PPM1a can also target and inhibit STAT1 in monocytes (27). Inhibition of PPM1a in monocytes leads to increased STAT1 activation. Therefore it is possible that the CLIC4 inhibits PPM1a in keratinocytes by binding to PPM1a and preventing it interacting with its substrates which would lead to elevated STAT1 activation.

In conclusion, the elevated type-1 IFN signalling found in the epidermis of SSc skin may in part be due to increased CLIC4 expression in the epidermis which leads to increased STAT1 activation and expression of ISGs. This is exciting from a therapeutic angle as targeting CLIC4 in SSc may prevent the inflammation associated with SSc and the fibrotic element of the disease (3,4). Therefore designing specific inhibitors that target CLIC4 ion channel function is an urgent need.

### Ethic Approval/IRB statement

The study was approved by NRES committee North East-Newcastle & North Tyneside: REC Ref:15/NE/0211 to FDG. For all in vivo mouse experiments there was strict adherence to institutional guidelines was practiced, and full local ethics committee approval through the UCL Animal Welfare and Ethical Review Body (AWERB) and appropriate Home Office Project and Personal license through CPD.

### Patient Consent Statement

All participants provided written informed consent to participate in this study. Informed consent procedure was approved by NRES-011NE to FDG by the University of Leeds.

### Patient and Public Involvement

Patients or the public were involved in the design, or conduct, or reporting, or dissemination plans of our research

## Data availability Statement

All data is included in the manuscript and accompanying supplementary material

## Author Contribution

Designing Research Study: C.W.W, F.D.G

Funding acquisition: C.W.W, F.D.G

Conducting Experiments: C.W.W, S.D, J.B, B.C-R, E.M.C, R.W, V.K, E.D.L, E.D.S

Acquiring Data: C.W.W, S.D, J.B, B.C-R, R.W, V.K, E.D.L, E.D.S

Analysing Data: C.W.W, R.L.R, C.P.D, P.J.M, F.D.G

Writing Manuscript: C.W.W, F.D.G

All authors reviewed and approved the final version of the manuscript

---

**Supplementary Figure 1:**
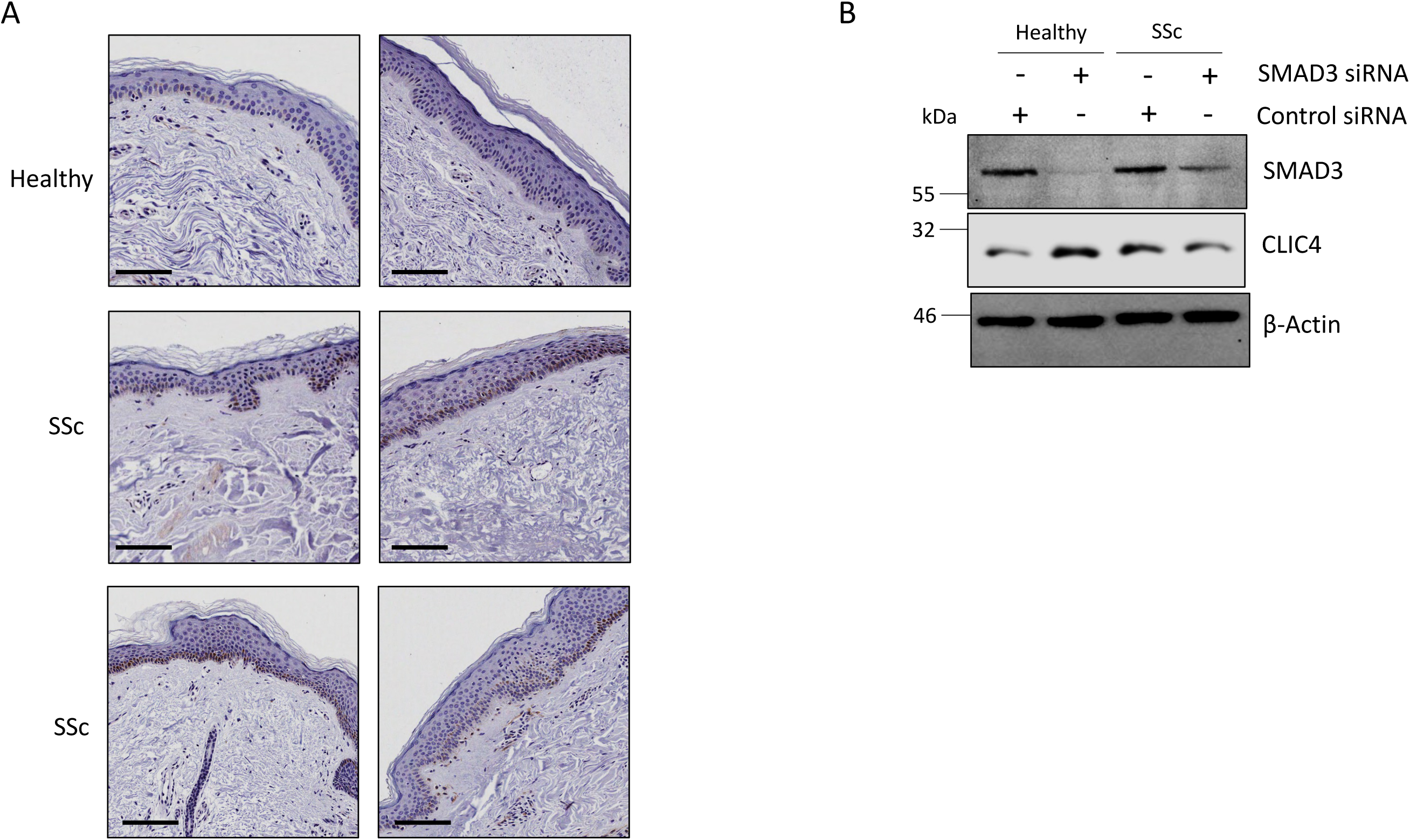
CLIC4 expression is upregulated in SSc skin keratinocytes. (A) Skin biopsies from healthy and SSc patient forearms were stained with an antibody specific to CLIC4 and visualized with an HRP conjugated secondary antibody. Scale bars represent 50μM. (B) Protein was extracted from healthy and SSc fibroblasts transfected with scramble control or SMAD3 siRNA for 72 hours. SMAD3 and CLIC4 protein levels were assessed by western blot. β-actin was used as a loading control.

**Supplementary Figure 2:**
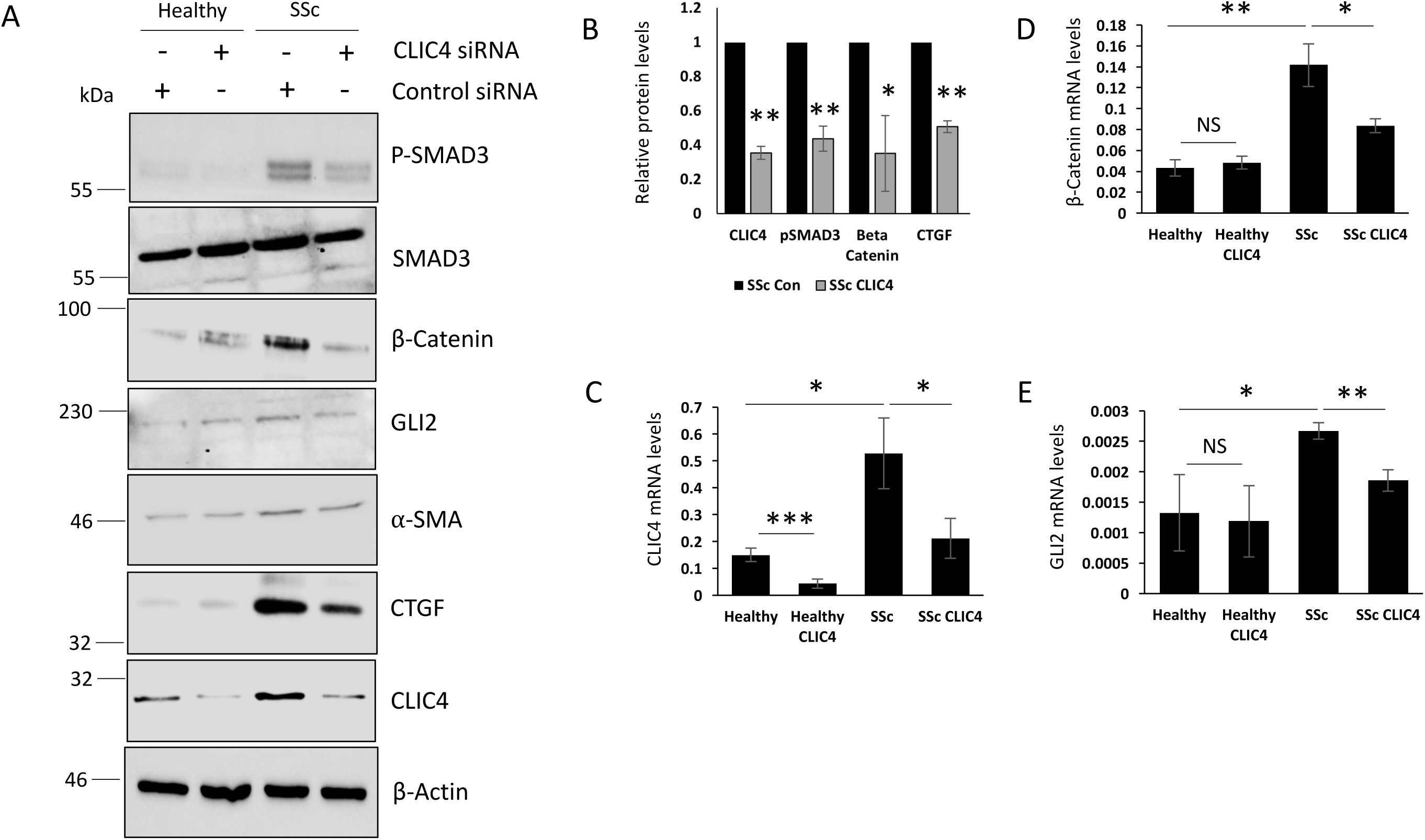
Knockdown of CLIC4 disrupts the pro-fibrotic phenotype in SSc dermal fibroblasts. Protein and RNA were extracted from healthy and SSc dermal fibroblasts transfected with siRNA specific for CLIC4. (A) pSMAD3, total SMAD3, β-catenin, GLI2, CTGF, α-SMA and CLIC4 protein levels were assessed by western blot. β-actin was used as a loading control. (B) Graph represent densitometry analysis for the mean and standard error for three independent experiments. CLIC4 (C), β-catenin (D) and GLI2 (E) transcript levels were assessed by qPCR.

**Supplementary Figure 3:**
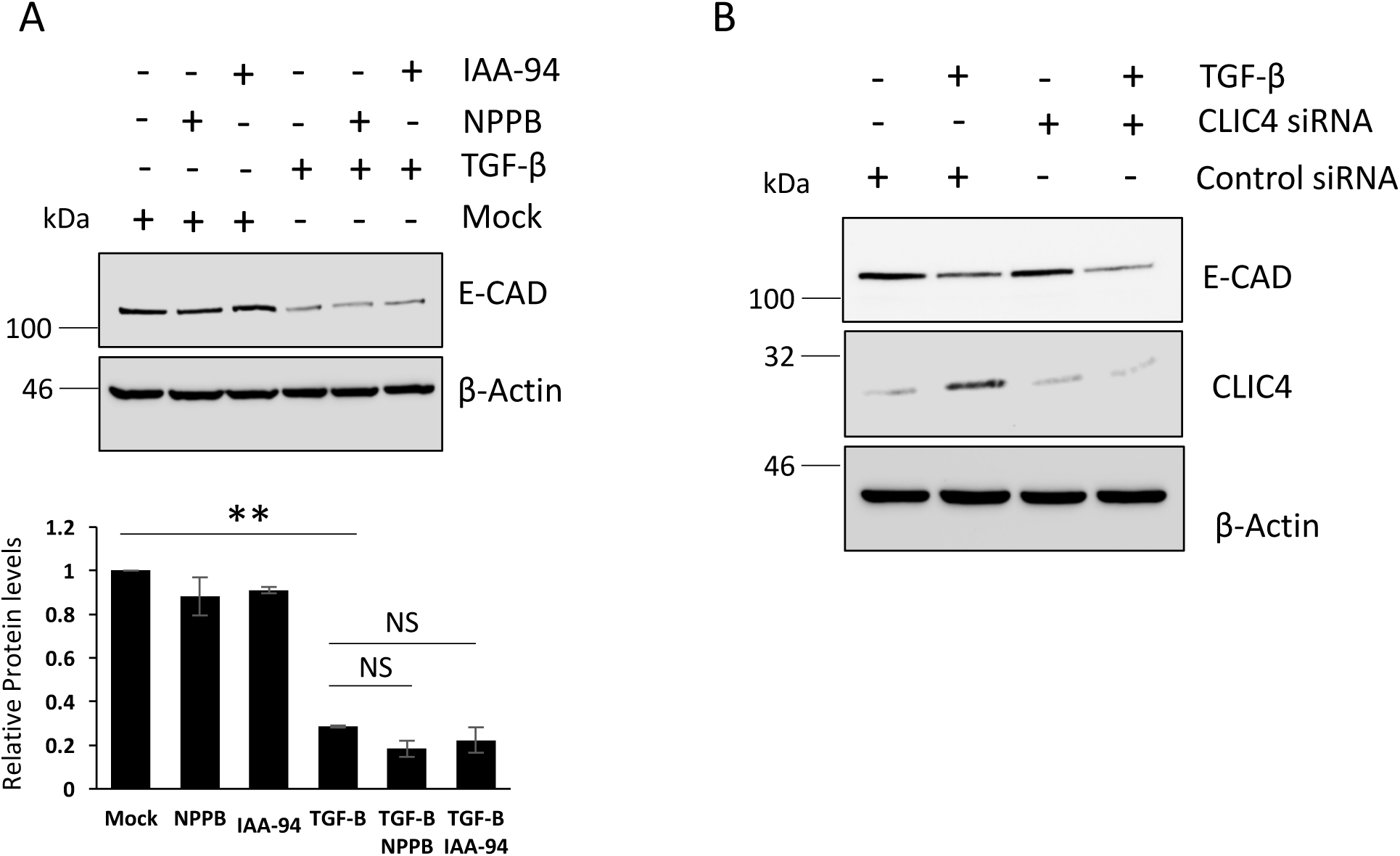
CLIC4 does not play a role in TGF-B associated EMT in keratinocytes. Protein was extracted from Hacats stimulated with TGF-β for 48 hours. In addition, Hacats were treated with the chloride channel inhibitors NPPB and IAA-94. (A) E-cadherin protein levels were assessed by western blot. β-actin was used as a loading control. Graphs represent densitometry analysis for the mean and standard error for three independent experiments. (B) Protein was extracted from, Hacats were transfected with the scramble and CLIC4 siRNA. After 24 hours the transfected Hacats were stimulated with TGF-β for a further 24 hours. E cadherin and CLIC4 protein levels were assessed by western blot. β-actin was used as a loading control.

**Supplementary Figure 4:**
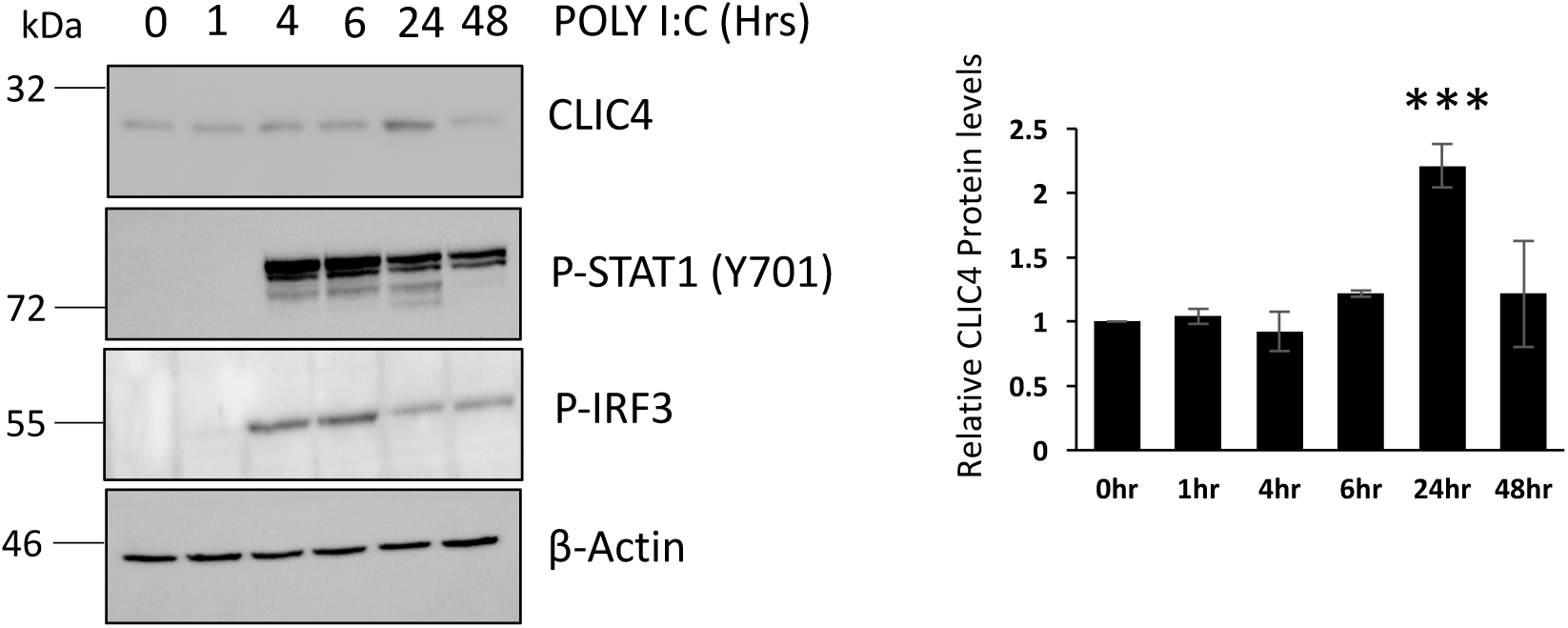
POLY I:C induces CLIC4 expression in a time dependent manner. (A) Hacat cells were stimulated with POLY I:C for between 1-48hrs. Protein was extracted from the cells and pSTAT1, CLIC4 and pIRF3 protein levels were analysed by western blot. β-actin was used as a loading control. Graphs represent densitometry analysis for the mean and standard error for three independent experiments.

**Supplementary Figure 5:**
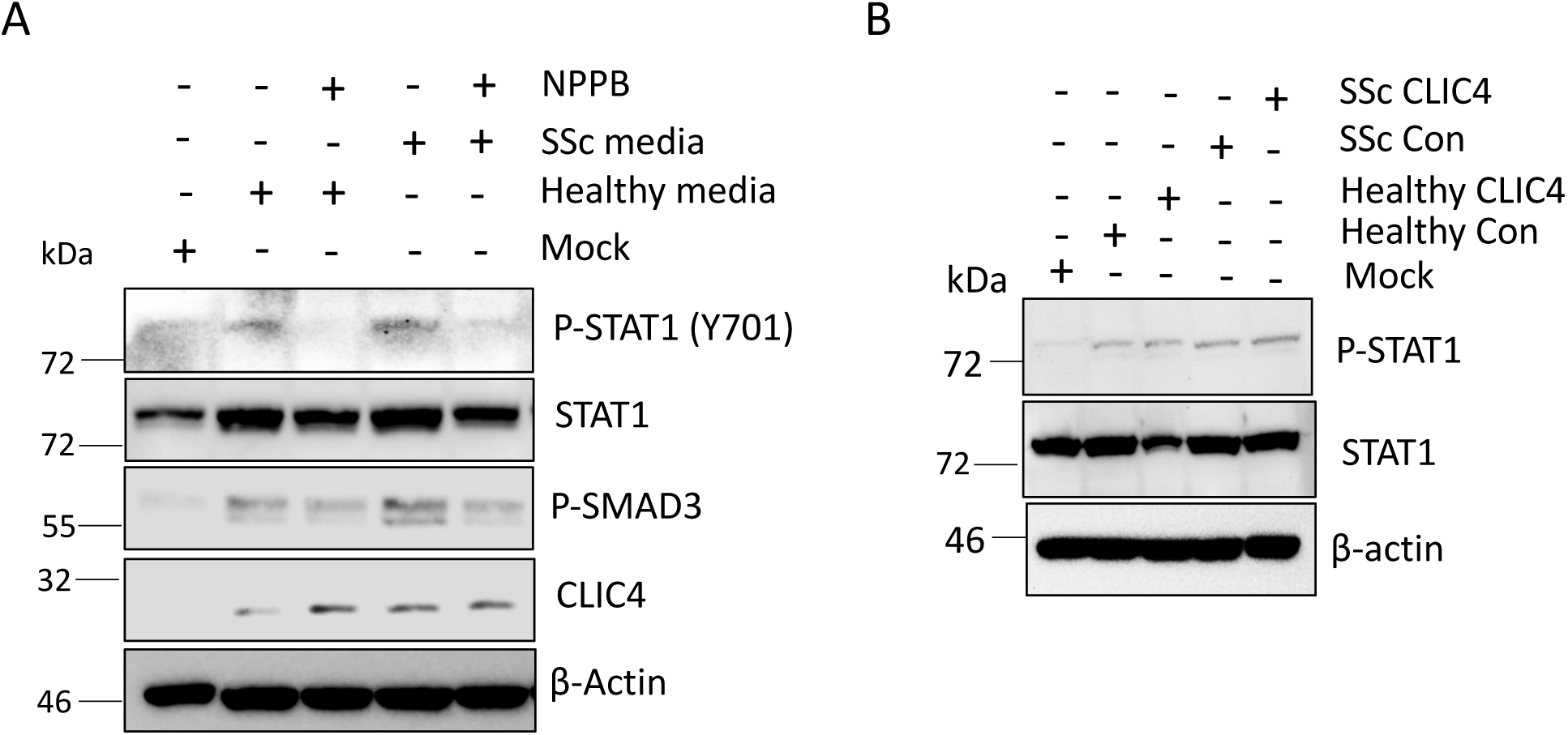
Inhibition of CLIC4 blocks SSc derived STAT1 activators. (A) Serum depleted conditioned media was collected from primary healthy and SSc patient fibroblasts after 48hrs. Hacats were stimulated with the media for 48 hours in the absence or presence of NPPB. pSTAT1, STAT1 pIRF3, pSMAD3 and CLIC4 protein levels were assessed by western blot. β-actin was used as a loading control. (B) Conditioned media was collected from healthy and SSc fibroblasts transfected with scramble and CLIC4 siRNA. The media was used to stimulate Hacats. pSTAT1 and β-actin protein levels were assessed by western blot.

**Supplementary Figure 6:**
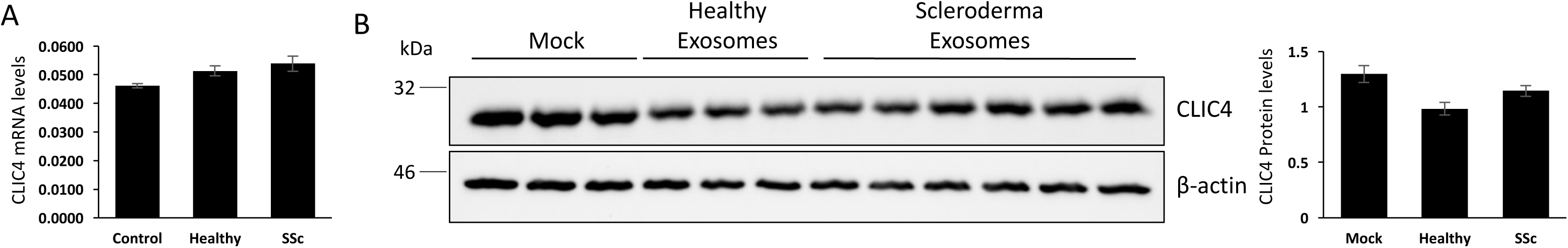
SSc fibroblast exosomes do not stimulate CLIC4 expression in keratinocytes. Exosomes isolated from healthy and SSc patient fibroblasts were used to stimulate Hacats for 48 hours. (A) RNA was isolated from stimulated hacats and CLIC4 transcript levels were assessed. (B) Protein was isolated from stimulated Hacats and CLIC4 protein levels were assessed by western blot. Graphs represent the mean and standard error for densitometry analysis.

