## Supplementary Materials and Methods for "Chloride intracellular channel 4 (CLIC4) is a global regulator of type 1 interferon signaling in Systemic Sclerosis (SSc) epithelial cells"

**Patient cell lines**

Full thickness skin biopsies were surgically obtained from the forearms of four adult healthy controls and four adult patients with recent onset SSc, defined as a disease duration of less than 18 months from the appearance of clinically detectable skin induration. All patients satisfied the 2013 ACR/EULAR criteria for the classification of SSc and had diffuse cutaneous clinical subset as defined by LeRoy et al (15). All participants provided written informed consent to participate in the study. Informed consent procedures were approved by NRES-011NE to FDG. Fibroblasts and keratinocytes were isolated and established as previously described (16). Primary cells were immortalized using human telomerase reverse transcriptase (hTERT) to produce healthy control hTERT and SSc hTERT.

**Cell culture**

hTERT patient fibroblasts, the keratinocyte cell line Hacats and lung epithelial cell line A549 were all maintained in Dulbecco’s modified Eagle medium (DMEM) (Gibco) supplemented with 10% FBS (Sigma) and penicillin-streptomycin (Sigma). Primary SSc keratinocytes were maintained in keratinocyte growth media (Promocell). Human umbilical vein endothelial cells (HUVECs) were maintained in ECGM2 media. Cells were treated with the chloride channel inhibitors (NPPB (25μM), IAA-94 (50μM)) or JAK1 inhibitor (Tofacitinib (2μM)) or TGF-B receptor inhibitor (SD-208 (1μM)) for 48 hours in a humidified incubator at 37C and 5% CO2.

**Trans-well Co-Culture Experiments**

Healthy and SSc fibroblasts were seeded onto 0.4-micron pore polyethylene terephthalate (PET) transmembranes (Corning). The transmembrane was inserted in wells containing an equal number of Hacats. After 48 hours, the well was removed and the Hacats were harvested for protein.

**Immune agonist stimulation**

Healthy dermal fibroblasts were serum starved for 24 h in DMEM containing 0.5% FBS and stimulated with Poly IC (10μg/ml), Poly dAdT (50ng/ml), IFNa (2ng/ml), ODN2216 (1μM) for 48 hours in combination with the Chloride channel inhibitors.

**siRNA transfections**

A pool of four siRNAs specific for different regions of CLIC4 or a negative control scrambled siRNA (Qiagen) were transfected into Hacats or A549 cells using Lipofectamine 2000 (Thermo Fisher). Hacats were incubated for 48 hours prior to harvesting and A549 cells incubated for 72 hours prior to harvesting. A pool of four siRNAs specific for different regions of SMAD3 or a negative control scrambled siRNA (Qiagen) were transfected into healthy and SSc dermal fibroblasts using Lipofectamine 2000. Sera depleted media was added to the fibroblasts after 24 hours and incubated for 48 hours prior to protein harvesting and media collection for downstream stimulation.

**Western blotting**

Total proteins were extracted from fibroblasts in RIPA buffer and resolved by SDS-PAGE (10-15% Tris-Glycine). Proteins were transferred onto Hybond nitrocellulose membranes (Amersham biosciences) and probed with antibodies specific for α-smooth muscle actin (Abcam), CLIC4 (Santa Cruz), phosphorylated and total STAT1 (Cell signalling), CTGF (Abcam), phosphorylated IRF3 (Abcam), β-catenin (cell signalling), phosho-SMAD3 (S423/S425) (Abcam), total SMAD3 (Cell signalling), E-Cadherin (Santa Cruz), CXCL10 (Abcam) and β-Actin (Sigma). Immunoblots were visualized with species-specific HRP conjugated secondary antibodies (Sigma) and ECL (Thermo/Pierce) on a Biorad chemiDoc imaging system.

**Quantitative Real time PCR**

RNA was extracted from cells using commercial RNA extraction kits (Zymo Research). RNA (1ug) was reverse transcribed using cDNA synthesis kits (Thermo). QRT-PCRs were performed using SyBr Green PCR kits on a Thermocycler with primers specific for MX1 (Forward: CGACACGAGTTCCACAAATG Reverse: AAGCCTGGCAGCTCTCTACC), CXCL10 (Forward; GGTGAGAAGAGATGTCTGAATCC Reverse; GTCCATCCTTGGAAGCACTGCA), CXCL11 (Forward; TCCCCCATGTTCAAAAGAGGAC Reverse; ATATCTGCCACTTTCACTGCTTTTAC), IFIT1 (Forward; GACTGGCAGAAGCCCAGACT Reverse; GCGGAAGGGATTTGAAAGCT), CLIC4 (Forward: CATCCGTTTTGACTTCAGTGTTG; Reverse: AGGAGTTGTATTTAGTGTGACGA) and GAPDH (Forward; ACCCACTCCTCCACCTTTGA Reverse; CTGTTGCTGTAGCCAAATTCGT). Data were analysed using the ΔΔ Ct method. GAPDH served as a housekeeping gene.

**Immunohistochemistry**

Immunohistochemistry was performed as previously described (1). Sections were stained with CLIC4 (1/200) (Santa Cruz) or Phosphorylated STAT1 (1/200) (cell signalling) antibodies, visualised using an HRP conjugated mouse secondary and counterstained with haematoxylin. Skin sections from age matched wildtype and TβRIIΔK-fib transgenic mouse model of human SSc were also stained for CLIC4 expression in additional control experiments.

**SSc fibroblast conditioned media stimulation**

Sub-confluent healthy and SSc dermal fibroblasts were grown in sera depleted DMEM for 48 hours. The media was collected and centrifuged at 2000g for 30 mins to remove cell debris. The media was added to Hacat cells neat for 48 hours or to HUVECs cells neat for 24 hours.

**Exosome isolation and stimulation**

Healthy and SSc dermal fibroblasts were grown in DMEM containing exosome depleted FBS until they reached confluence. The media was collected and ultracentrifuged (42,000rpm, 2 hours). The exosome pellet was extracted using the total exosome isolate reagent (Invitrogen). Hacats were stimulated with 1% total volume of exosomes for 48 hours.
